# Reanalysis of global proteomic and phosphoproteomic data identified a large number of glycopeptides

**DOI:** 10.1101/233247

**Authors:** Yingwei Hu, Punit Shah, David J. Clark, Minghui Ao, Hui Zhang

## Abstract

Protein glycosylation plays fundamental roles in many cellular processes, and previous reports have shown dysregulation to be associated with several human diseases, including diabetes, cancer, and neurodegenerative disorders. Despite the vital role of glycosylation for proper protein function, the analysis of glycoproteins has been lagged behind to other protein modifications. In this study, we describe the re-analysis of global proteomic data from breast cancer xenograft tissues using recently developed software package GPQuest 2.0, revealing a large number of previously unidentified *N*-linked glycopeptides. More importantly, we found that using immobilized metal affinity chromatography (IMAC) technology for the enrichment of phosphopeptides had co-enriched a substantial number of sialoglycopeptides, allowing for a large-scale analysis of sialoglycopeptides in conjunction with the analysis of phosphopeptides. Collectively, combined MS/MS analyses of global proteomic and phosphoproteomic datasets resulted in the identification of 6,724 N-linked glycopeptides from 617 glycoproteins derived from two breast cancer xenograft tissues. Next, we utilized GPQuest for the re-analysis of global and phosphoproteomic data generated from 108 human breast cancer tissues that were previously analyzed by Clinical Proteomic Analysis Consortium (CPTAC). Reanalysis of the CPTAC dataset resulted in the identification of 2,683 glycopeptides from the global proteomic data set and 4,554 glycopeptides from phosphoproteomic data set, respectively. Together, 11,292 N-linked glycopeptides corresponding to 1,731 N-linked glycosites from 883 human glycoproteins were identified from the two data sets. This analysis revealed an extensive number of glycopeptides hidden in the global and enriched in IMAC-based phosphopeptide-enriched proteomic data, information which would have remained unknown from the original study otherwise. The reanalysis described herein can be readily applied to identify glycopeptides from already existing data sets, providing insight into many important facets of protein glycosylation in different biological, physiological, and pathological processes.

## Introduction

Protein glycosylation is one of the most common protein modifications universally present in all living organisms^1^. Glycosylation has diverse biological functions and plays critical roles in various biological activities, especially in cell recognition and adhesion^1–7^. With the development of high-throughput mass spectrometry-based proteomics, it is allowed for the identification and quantification of thousands of proteins and protein modifications for a comprehensive profiling of cells, tissues or body fluids^8–14^.

Analysis of the glycoproteome is challenging. Due to the complicated nature of glycosylation, wherein a glycoprotein could have multiple sites of glycosylation, with each site displaying heterogeneity due to variability in the attached glycan structure, most studies of glycoproteins are limited in exploring either the glycosite^15^ or released glycan structure separately^7,16,17^. However, information related to both glycosites and the attached glycans at specific glycosylation sites need be determined. More recently, analyses of intact glycopeptides have been used to study glycosylation heterogeneity from a specific glycoprotein or complex samples^18–24^ Recently, several software tools^25^ have been developed to assist the assignment of glycopeptides, such as Byonic^26^, Glycopeptide Search (GPS)^27^, Glycopep Grader^28^, SpectraST^29^, GPQuest^30^, and pGlyco^31,32^, allowing for high throughput identification of intact glycopeptides.

Recently, it has been reported that unaccounted mass spectrometry spectrum exists in proteomic data sets, and may be related to unidentified post-translational modifications^33^. With the preponderance of glycosylation as a common protein modification, we sought to explore the incidence of glycopeptides in non-glycocentric data sets. In this study, we re-analyzed data from global proteomics and phosphoproteomics using GPQuest 2.0 software tool to elucidate the presence of *N*-linked glycopeptides. In our analysis, we showed the existence of tens of thousands of glycopeptide spectra in non-glycoenriched data sets, yielding the identification of thousands of *N*-linked glycopeptides using GPQuest 2.0. Overall, these results further support the hypothesis of glycosylation as one of the most abundant modifications of proteins in human cells, and that glycopeptides are widely present in global and IMAC-enriched phosphorylated proteomic mass spectrometry data sets.

## Results

### The abundance of glycopeptides identified from the global and phosphoproteomic analyses of breast cancer xenograft tissues

To determine the abundance of glycopeptides in proteomic analysis, two human-in-mouse xenograft tissues were analyzed by proteomics and phosphoproteomics. This pair of xenograft tissues representing basal (P32) and luminal-B (P33) human breast cancer were used as reference materials to determine the longitudinal reproducibility of LC-MS/MS experiments during the analysis of cancer tissues in Clinical Proteomic Tumor Analysis Consortium (CPTAC) project^8,11–14^ The pair of xenograft tissues were tryptic digested, labeled by TMT, fractionated by bRPLC, and 5% of fractionated peptides were analyzed by LC-MS/MS for global proteomics. The remaining 95% of peptides were used for the enrichment of phosphopeptides using IMAC and analyzed by LC-MS/MS. We first utilized the database search tool MSGF+^34^ to identify peptides without modifications. As shown in Figure 1A and Supplementary Table 1, MSGF+ assigned over 27% of total 1,163,831 HCD MS/MS spectra from the 24 fractions of global proteomics data, with FDR <= 1%. We then determine whether the un-assigned spectra could come from glycopeptides. It is well known that the HCD fragmented oxonium ions in MS/MS spectra are reliable indicators of glycopeptides^19^. The spectra containing HexNAc oxonium ion (m/z 204.0966) in the top 10 most intense peaks are regarded as potential candidates of the glycopeptide-related spectra and named as ‘oxo-spectra’. The search results of oxo-spectra in the global proteomics data sets are highlighted as yellow in Figure 1A, revealing that approximately 1% MS/MS spectra (12,896) were identified as ‘oxo-spectra’.We employed the same search method to explore the phosphoproteomics data set, with the results shown in Figure 1B. MSGF+ assigned approximately 12.1% of the total 590,995 HCD MS/MS spectra from 13 fractions of phosphoproteomics data to phosphopeptides with FDR <= 1%. Of note, we found more than 30% of the MS/MS spectra identified as ‘oxo-spectra’ in the phosphoproteomic data set. These results show that there was approximately about 30 times more potential glycopeptide-related spectra observed in phosphoproteomics dataset than in the global proteomics dataset. Surprisingly, we observed the number of ‘oxo-spectra’ was higher than the number of identified spectra assigned as phosphopeptides by MSGF+ in the phosphoproteomics dataset, indicating that the IMAC-based phosphopeptide affinity capture could also enrich glycosylated peptides. Taken together, these results showed that intact glycopeptides are widely present in these global and phosphoproteome datasets from human-in-mouse xenograft tissues.

**Figure 1.**
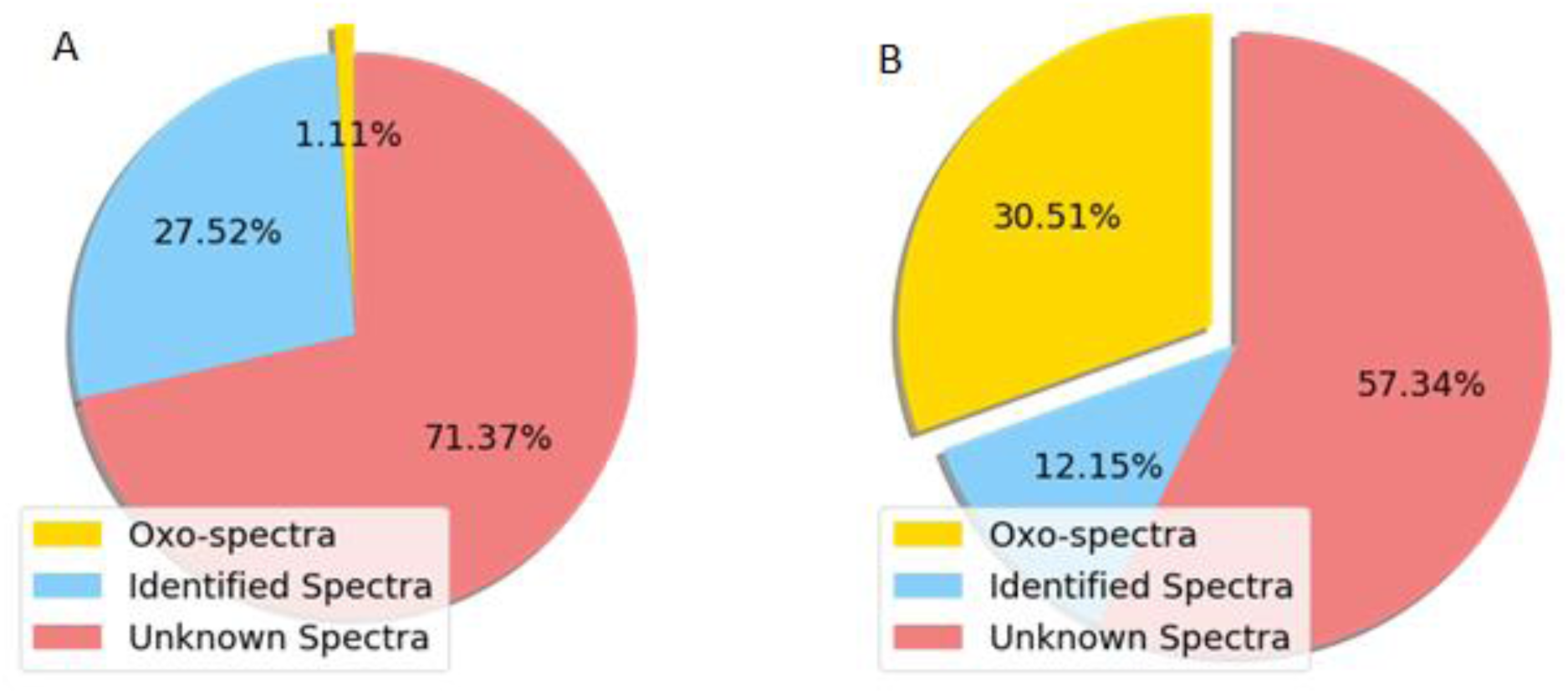
The preliminary estimation of potential glycopeptides in the global and phosphoproteomic data sets from breast cancer xenograft tissues. The search results of global and phosphopeptide-enriched datasets are shown in A and B pie charts respectively. The blue chart represents the proportion of identified peptides (A) and phosphopeptides (B). The red part presents the proportion of unidentified spectra. The yellow part represents the proportion of ‘oxo-spectra’.

### Identification of N-linked glycopeptides of the breast tumor xenograft by GPQuest

To facilitate identification of intact N-linked glycopeptides from resulting ‘oxo-spectra’, the global and IMAC-based phosphopeptide-enriched phosphoproteomics data sets of the breast cancer xenograft tissues were searched using GPQuest against a human database containing over 30,000 known N-linked glycopeptide sequences and 181 *N*-linked glycan compositions^35^. The search results are summarized in Table 1. In the global proteomics data set, there were 2,631 MS/MS spectra assigned as intact *N*-linked glycopeptides by GPQuest with FDR <= 1%, corresponding to 1,660 *N*-linked glycopeptides, 427 *N*-linked glycosites and 111 N-linked glycan compositions from 266 *N*-linked glycoproteins. Over 20% ‘oxo-spectra’ (2,631/12,896) were successfully assigned as *N*-linked glycopeptides. In the phosphoproteomics data, there were 9,836 MS/MS spectra assigned as intact *N*-linked glycopeptides by GPQuest with FDR <= 1%, corresponding to 5,987 *N*-linked glycopeptides, 1,041 *N*-linked glycosites and 163 *N*-linked glycan compositions from 589 *N*-linked glycoproteins (See Table 1 and Supplementary Table 2). It is not surprising that there were more *N*-linked glycopeptides identified from the phosphoproteomics data relative to the global proteomics data based on the glycopeptide abundances estimated by the number of oxo-spectra (see Figure 1).

**Table 1.** Summary of glycoproteomic results from GPQuest search of Xenograft and CPTAC data sets

|  | *Xenograft*<br>*Global* | *Xenograft*<br>*Phospho* | *CPTAC*<br>*Global* | *CPTAC*<br>*Phospho* |
| --- | --- | --- | --- | --- |
| *PSM* | 2631 | 9836 | 26994 | 44865 |
| *Glycopeptide* | 1660 | 5987 | 2683 | 4554 |
| *Glycosite* | 427 | 1041 | 727 | 978 |
| *Glycoprotein* | 266 | 589 | 420 | 514 |
| *Glycan* | 111 | 163 | 114 | 144 |

### Sialylated glycopeptides were enriched in phosphoproteomics dataset

The compositions of identified glycans in both global and IMAC-based phosphopeptide-enriched proteomics data were further analyzed to determine the type of glycans identified in these two proteomics datasets. We observed a higher percentage of glycans containing sialic acid (28%) in the phosphoproteomics data compared to that of the global proteomics data (9%), suggesting that the IMAC-based phosphopeptide enrichment method may selectively enrich sialylated glycans (See Figure 2A). To determine the glycans specifically enriched in phosphoproteomics dataset, we investigated the distribution of PSMs of different glycan types. The PSMs were grouped by the sample and the glycan type attached to the assigned *N*-linked glycopeptides. For classification of intact glycopeptides, glycans containing HexNAc=2, Hex >=5 and no other monosaccharides were classified as ‘Oligomannose’, glycans containing HexNAc >=2, Hex >=3, Sialic Acid >=1 grouped as ‘Sialylated’, and finally all remaining glycans assigned in the ‘Others’ group, which included non-sialylated hybrid or complex glycans. As shown in Figure 2B, the phosphoproteomics data were found to be highly enriched for sialylated *N*-linked glycopeptides compared to global proteomics datasets. We observed a less enrichment for non-sialylated hybrid or complex glycans, or oligomannose N-linked glycopeptides, which indicated that IMAC enrichment for phosphopeptides displayed a co-enrichment for sialylated glycopeptides and less enrichment for glycopeptides with neutral glycans.

**Figure 2.**
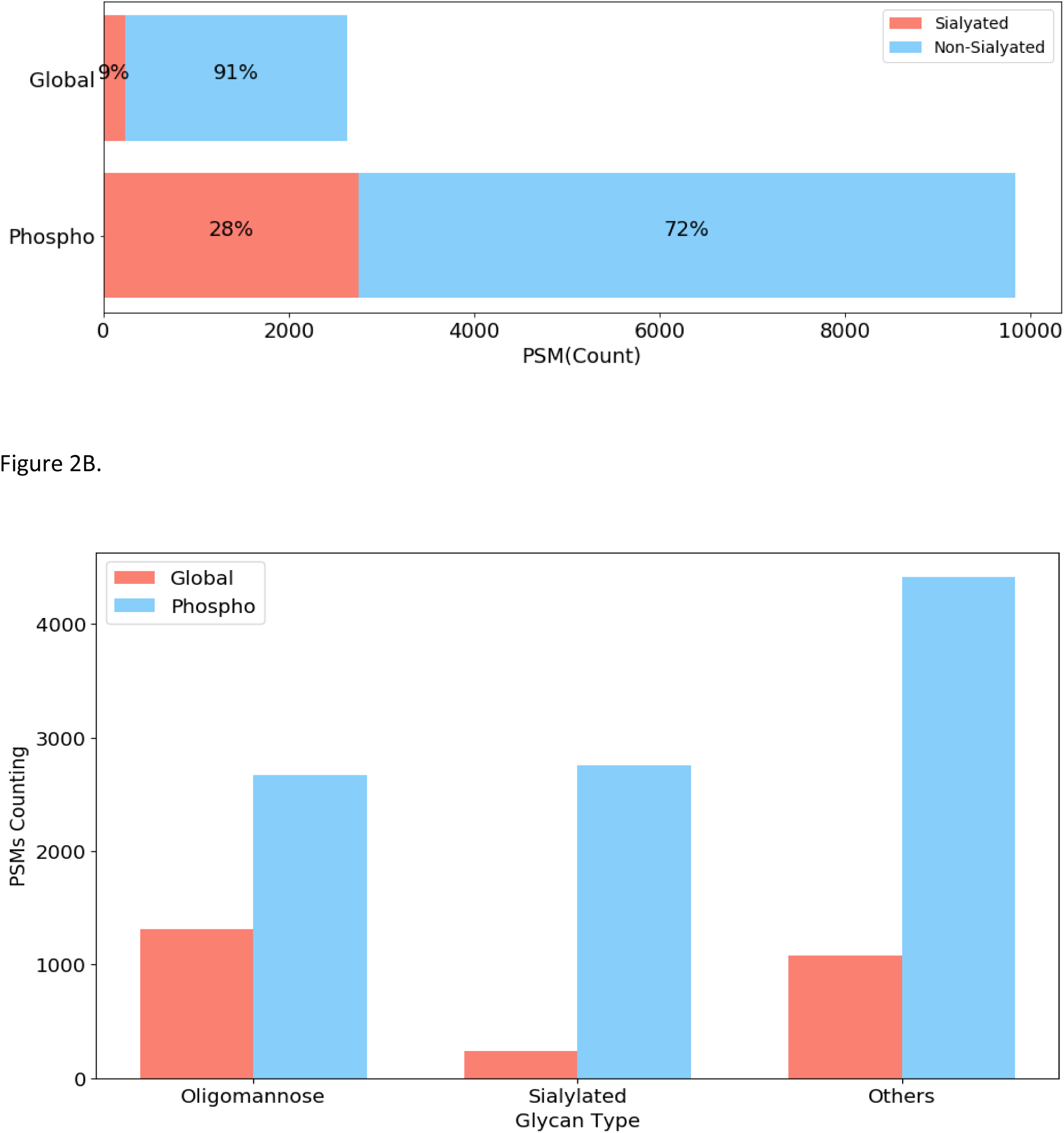
The classified glycopeptide-spectrum matchings (PSMs) of different glycopeptides from global proteomics and IMAC-enriched phosphoproteomics datasets of breast cancer xenograft tissues. A. The red bars are the percentage of PSMs assigned as N-linked sialoglycopeptides. The blue bars are the percentage of PSMs from non-sialylated glycopeptides. B. The total number of glycopeptide-spectrum matchings (PSMs) of different glycan types in global proteomics and IMAC-enriched phosphoproteomics datasets.

### Intact glycopeptide analysis on CPTAC breast cancer tumors

For comprehensive proteome and phosphoproteomic analyses of breast cancer, a total of 108 breast tumors, were analyzed with a total of 72 iTRAQ datasets (36 global proteomics and 36 IMAC-enriched phosphoproteomics datasets)^11^. The datasets were analyzed using MSGF+, “oxo-spectra” and GPQuest as described in the analysis of breast xenograft tissues. On average, We observed 4,857 ‘oxo-spectra’ in each global proteomics dataset (Supplementary Table 3 and Figure 3), whereas the average number of PSMs of phosphopeptides identified by MSGF+ in each global data set was 3,617. This result suggests that glycopeptides in global samples without any enrichment are abundant, the total number of glycan containing spectra (‘oxo-spectra’) is similar to the number of the spectra identified as phosphopeptides, which is currently considered as one of the most abundant protein modifications in eukaryotes^36^. The median number of intact *N*-linked glycopeptides, glycosites, glycans, and glycoproteins identified by GPQuest with FDR <= 1% in each global proteomics dataset was about 527, 238, 61, and 164, respectively. In sum, 26,994 PSMs, 2,683 *N*-linked glycopeptides from 727 glycosites, 420 glycoproteins and 114 glycans were identified from 36 global proteomic datasets. In each phosphoproteomics dataset, there were averagely 23,100 ‘oxo-spectra’. The median number of *N*-linked glycopeptides, glycosites, glycans, and glycoproteins identified by GPQuest with FDR<= 1% (glycopeptide-spectrum matching GPSM level) in each phosphoproteomics dataset was around 611, 247, 69, and 163 respectively. In sum, 44,865 PSMs, 4,554 *N*-linked glycopeptides from 978 glycosites, 514 glycoproteins, and 144 glycans were identified from 36 phosphoproteomic datasets (Supplementary Table 4).

**Figure 3.**
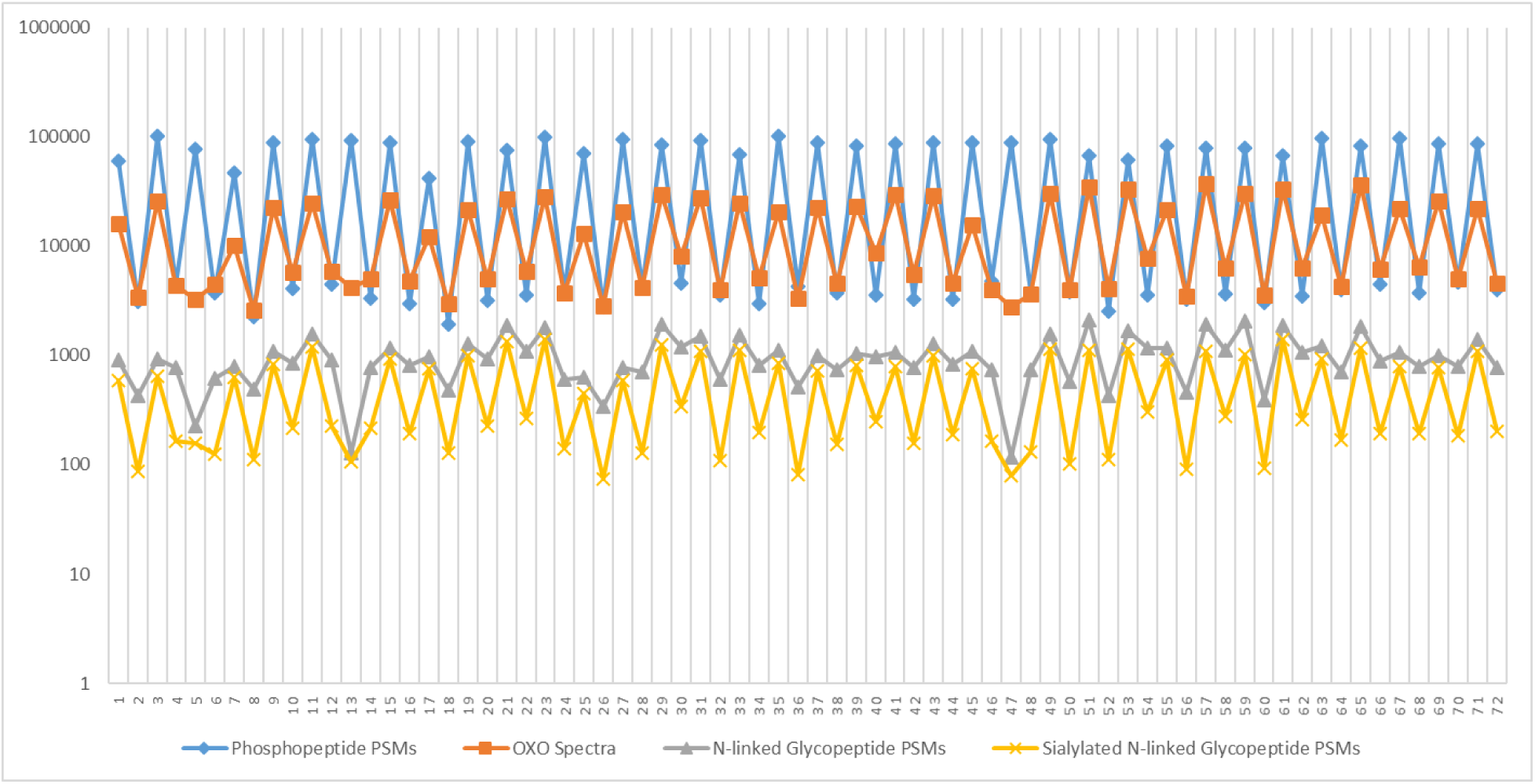
The abundance of phosphopeptides and glycopeptides in all the samples of ‘Breast Cancer CPTAC’ data. All the 72 data sets (36 global and 36 phosphopeptide-enriched datasets) of the CPTAC breast cancer cells were searched for phosphorylation (Blue), oxo-spectra (Red), intact N-linked glycopeptides (Grey) and sialylated N-linked glycopeptides (Yellow). The results of PSMs/spectral counting were plotted together. The x-axis is the sample index. All the odd index numbers represent phosphoproteomic data sets. All the even index numbers represent the global proteomic data sets of the sample with an index of N-1, where N is the even index number. The Y-axis is the log-scaled PSMs/spectral counts.

The number of assigned phosphopeptides and *N*-linked glycopeptides from all 72 datasets are summarized in Figure 3. We plotted the log (PSM/spectral count) of identified phosphopeptides, ‘oxo-spectra’, N-linked glycopeptides, and sialylated N-linked glycopeptides against the phosphoproteomics datasets (odd sample ID number) and the corresponding global proteomics datasets from the identical breast tumors (even sample ID number).The results showed the abundance of ‘oxo-spectra’ and N-linked glycopeptides, especially sialylated N-linked glycopeptides were strongly positively correlated to the spectral counting of phosphopeptides in nearly all the datasets except 3 cases (No. 5,13 and 47), which illustrated that the IMAC-based phosphopeptide affinity capture method co-enriched glycopeptides from the original samples in most of the cases. In total, 71,859 Glycopeptide-Spectrum Matchings (GPSMs), 5,625 *N*-linked glycopeptides, 1,163 glycosites and 146 glycans were identified from 590 glycoproteins of the 72 datasets. The complete results of all datasets are shown in Supplementary Table 4.

## Discussion

Glycosylation is one of the most abundant and important modifications of protein. However, the identification of glycosylation is often omitted in the previous studies due to a multitude of factors, including their low abundance, dynamic expression and lack of complimentary approaches to investigate multiple protein modifications in parallel. However, with the rapid advances of mass spectrometry technologies, sample preparation, and novel search algorithm strategies, there has been a growing interest in analyzing a wide array of protein/peptide modifications including intact glycopeptides. In this study, we investigated the presence of *N*-linked glycoproteins in the global and IMAC-based phosphopeptide-enriched proteomics datasets of the breast cancer xenograft and CPTAC breast cancer samples, revealing a large number of *N*-linked glycopeptides in a total of 84,326 GPSMs, 11,292 *N*-linked glycopeptides, 1,731 glycosites, and 166 glycans identified from 883 glycoproteins that were omitted in the original studies. This observation has provided further evidence that glycosylation is one of the most abundant protein modifications, illustrating it’s significant role in biology and the importance of its inclusion in future analyses.

The development of glyco-specific algorithms and tools to interrogate MS/MS spectra is critical to identify glycopeptides. GlycopeptideSearch (GPS) is a semi-automated tool for the identification of *N*-linked glycopeptides^27^. It can generate a list of most likely candidates of the combinations of peptides and glycans but requires additional manual validation to report and ascertain a false discovery rate (FDR). GlycoPep Grader is a web-based utility for *N*-linked glycopeptide identification, which considers fragments from complexity composition of glycan structures to achieve more accurate glycan structure assignment^28^. It doesn’t support high-throughput searching. Another software tool, pGlyco, can be utilized for intact *N*-linked glycopeptide analysis with an FDR estimation^31,32^. The most recent version eliminated the dependency on the multiple fragmentation strategies (i.e. combining MS3 and HCD/ETD spectra), instead, required high quality of HCD MS/MS spectra using a stepped fragmentation method. Byonic is a commercial software tool for high-throughput intact glycopeptide analysis ^26^. Finally, the GPQuest used in this study is a freely available software focused on the high-throughput intact glycopeptide analysis, utilizing HCD MS/MS spectra^37^. The software provides FDR estimation for each assignment of the identified glycopeptide, as well as predicting glycosylation types based on generated oxonium ions^38^.

The presence of unknown spectra existing in shotgun proteomics experiment has been observed previously, even with proper search parameter settings^39^. The missed assignment of glycopeptides is one potential contributor to the unassigned spectra, resulting from the limitation of protein database search space or with defined protein modificatons^40^. With knowledge of oxonium ions indicating the fragmentation of a glycan structure, we can exploit this occurrence and search for ‘oxo-spectra’ that can then be used to roughly estimate the potential expression of glycopeptides in proteomic datasets. As a proof-of-concept, we showed that ‘oxo-spectra’ occupied about 1% of a global proteomics dataset and over one-third of the unassigned MS/MS spectra from an IMAC-based phosphopeptide-enriched proteomics dataset (Figure 1). The percentages of the ‘oxo-spectra’ indicated that adding glycopeptides to the search space is essential, especially in the IMAC-based phosphopeptide-enriched samples. In fact, thousands of *N*-linked glycopeptides were identified from the global proteomics and IMAC-enriched phosphoproteomics datasets as shown in Table 1, which is sufficient to be regarded as a large-scale profiling of protein *N*-linked glycosylation of breast cancer samples. Additional unassigned oxo-spectra in this study could be contributed by glycosylation other than *N*-linked glycosylation, which were not evaluated in this study for glycopeptide identifications. With increased understanding of spectral patterns from different glycosylation and further development of GPQuest software tool to include algorithms for other types of glycosylation, there should be additional glycopeptides identified from the ‘oxo-spectra’ that were not identified as N-linked glycopeptides.

According to the search results of ‘oxo-spectra’ and intact *N*-linked glycopeptides by GPQuest, phosphopeptide affinity capture using IMAC enrichment technology enriches not only phosphopeptides but also sialoglycopeptides, possibly due to the negative charges carried by both phosphopeptides and sialoglycopeptides. As shown in Figure 2A, 28% of the intact N-linked glycopeptides identified in the IMAC-based phosphopeptide-enriched samples contained a sialic acid residue but only 9% of those in global samples contain sialic acids. It is reasonable to believe that phosphopeptide affinity capture using IMAC enrichment technology may selectively enrich sialylated glycosylated peptides compared to other intact glycopeptides. Moreover, the identification of 72% glycopeptides without sialic acids could be related to the loss of sialic acids during the LC-MS/MS analysis after initial IMAC enrichment procedure, due to the labile nature of sialic residues during ionization^41^. This latter prospect is supported by the specific enrichment of hybrid and complex glycopeptides, but not glycopeptides with neutral oligomannose which would not contain the sialic residue. Further investigation of the influence of IMAC-enrichment on intact glycopeptides, specifically sialylated intact glycopeptides is warranted, as well as the scheme of the co-enrichment of both phosphopeptides and intact glycopeptides.

## Conclusions

With the rapid development of mass spectrometry and computational glycoproteomics tools, we can conduct a large-scale analysis of glycoproteome without specific glycopeptide enrichment and mass spectrometry analysis. In this study, we used our recently developed and improved intact glycopeptide analysis tool, GPQuest 2.0, to investigate the glycopeptide expression in two large datasets: two breast cancer xenograft samples and 108 CPTAC breast cancer tissues. The search results of ‘oxo-spectra’ and intact *N*-linked glycopeptides analysis demonstrated the feasibility of profiling glycoproteome utilizing global proteomics or the phosphoproteomics datasets. It was also shown that the IMAC-based phosphopeptide enrichment technology has a specific preference for glycopeptides containing sialylated glycans. There is no doubt that the information of glycoproteins from the additional intact glycopeptide analysis can provide a new dimension of in viewing the glycoproteome, as well as help us further understand the complicated biological role of glycosylation in nature.

## Methods

### Sample preparation of ‘Breast Cancer Xenograft’ sample

The xenograft tumor samples acquired from CPTAC program were generated from primary or metastatic breast tumors^11–14^. The tumor samples were lysed with sonication in 8M Urea and 1M NH4HCO3 pH8, containing 75mM NaCl. Inhibitors of phosphatase and O-GlcNAcase were added in the lysis buffer. After lysis, proteins were reduced with 5mM DTT, alkylated with 10mM IAA and digested with LysC and trypsin (Promega) at 37°C. The digested peptides were desalted on C18 SepPak columns (Waters). 400 μg desalted peptides were then labeled by an individual channel of TMT10plex (Thermo Fisher Scientific). After TMT label reaction, all ten channels were combined and desalted on a C18 SepPak column. Basic reversed phase fractionation was performed on Agilent 1100 HPLC analytical system, generating 24 fractions for global proteome analysis and 13 combined fractions for phosphopeptide enrichment. Phosphopeptides were enriched from each of the 13 fractions using IMAC method and desalted by stage-tips.

### Mass spectrometry analysis

Each fraction of global proteomics data and phosphoproteomics data was analyzed on a Lumos instrument (Thermo Fisher Scientific) once with Data-Dependent Acquisition (DDA). The DDA run consisted of one MS survey scan (60,000 resolution; AGC target: 4.0e5; mass range: 350-1800 m/z; charge state include: 2-6) followed by 20 MS/MS scans (15,000 resolution; AGC target: 5.0e4; HCD collision energy: 37 eV), with former precursors excluded for 45s after being selected once (dynamic exclusion option).

### Datasets

The ‘Xenograft’ dataset of the breast cancer cells generated above was called as ‘Breast Cancer Xenograft’ or ‘Xenograft’ dataset in this study. The other datasets of CPTAC breast cancer cells were downloaded from the CPTAC Data Portal^11^ (https://cptac-data-portal.georgetown.edu/cptacPublic/). The whole data repository was generated for proteogenomic analysis of TCGA breast cancer samples. It contains 72 datasets of 108 TCGA samples (36 global proteomic data sets and 36 phosphoproteomic data sets). IMAC-enriched phosphoproteomics and global proteomics datasets were marked in odd and even dataset ID numbers respectively. There were 13 fractions of each phosphopeptide-enriched dataset and 25 fractions of each global proteomics dataset in each group of 3 TCGA samples and one pooled reference sample. All the mzML files of the 72 datasets were downloaded and studied as ‘Breast Cancer CPTAC’ or ‘CPTAC’ dataset in the following analysis.

### Searching ‘oxo-spectra’ and *N*-linked glycopeptides by GPQuest 2.0

The RAW files from the Lumos instrument were converted to .mzML format by ProteoWizard package with the ‘Peak Picking’ option selected for all MS levels. GPQuest 2.0 was applied to investigate the expression of protein glycosylation on the unidentified MS/MS spectra in two approaches: the searching of spectra containing oxonium ions (‘oxo-spectra’) and the identification of intact *N*-linked glycopeptides. The oxonium ions were regarded as the signature features of the glycopeptides assigned to the MS/MS spectra, which were caused by the fragmentation of glycans attached to intact glycopeptides in the mass spectrometer. In this study, the MS/MS spectra containing the oxonium ions (m/z 204.0966) in the top 10 abundant peaks after removing reporter ions (TMT or iTRAQ) were considered as the potential glycopeptide candidates and named as ‘oxo-spectra’. The number of ‘oxo-spectra’ was used as a preliminary estimation of protein glycosylation expression. The intact *N*-linked glycopeptides were identified by using GPQuest to search against a customized database containing over 30,000 known *N*-linked glycopeptide sequences of human cells and a database containing 181 *N*-linked glycan compositions. The database of *N*-linked glycopeptides was collected from the previous literature and unpublished results in our lab. The glycan database was collected from the public database of GlycomeDB^42^(http://www.glycome-db.org). The theoretical b/y ions fragments of all the target and decoyed candidates of glycopeptides were calculated and built as a fragment ion index similar in MSFragger^33^ but applied in a different searching strategy without precursor mass restriction. The binned m/z values of fragments and their belonged peptides were stored as <key,value> pairs in a hashed map as the search space. Each tandem mass spectrum was first processed in a series of preprocessing procedures, including removing reporter ions (TMT or iTRAQ), spectrum denoising, intensity transformation, oxonium ions evaluation and glycan types prediction. The top 100 peaks in each qualified preprocessed spectrum were matched to the fragment ion index to find all the candidate peptides. The candidate peptides of each peak were merged and sorted descendingly according to their number of shared peaks. If the number of shared peaks is less than a threshold ( 6 peaks in this study), these candidate peptides were filtered. All the remaining candidate peptides were compared with the spectrum again to calculate the Morpheus scores^43^ by considering all the peptide fragment, glycopeptide fragments, and their isotope peaks. The peptide having highest Morpheus score was assigned to the spectrum finally. The mass gap between the assigned peptide and the precursor mass was searched in the glycan database to find the associated glycan. The best hits of all ‘oxo-spectra’ were ranked by the Morpheus score descendingly, in which those under FDR=1% and covering over 20% total intensity of the spectrum were reserved as qualified identifications. In this strategy, no precursor mass or Y ion was required in searching the peptide which decreases the loss of identification due to incorrect precursor mass or charge assignment. The precursor mass tolerance was set as 10ppm, and the fragment mass tolerance was 20 ppm.

### Search phosphopeptides by MSGF+

To compare the abundances of glycopeptides and phosphopeptides in both global proteomics and IMAC-enriched phosphoproteomics datasets, MSGF+ was applied to perform the searching of phosphopeptides. MSGF+ is a commonly used search engine in the conventional proteomics approach. In the searching of Xenograft samples of breast cancer, the protein database contains 32,799 proteins of human and 32,800 proteins of mouse. The missed cleavage is 2. The *in silico* cleavage was performed by using trypsin rule. All the other parameters were set by default.

